# Evidence of a pan-tissue decline in stemness during human aging

**DOI:** 10.1101/2023.04.13.536766

**Authors:** Gabriel Arantes dos Santos, Gustavo Daniel Vega Magdaleno, João Pedro de Magalhães

## Abstract

Despite their biological importance, the role of stem cells in human aging remains to be elucidated. In this work, we applied a machine learning methodology to GTEx transcriptome data and assigned stemness scores to 17,382 healthy samples from 30 human tissues aged between 20 and 79 years. We found that ∼60% of the studied tissues present a significant negative correlation between the subject’s age and stemness score. The only significant exception to this pattern was the uterus, where we observed an increased stemness with age. Moreover, we observed a global trend of positive correlations between cell proliferation and stemness. When analyzing the tissues individually, we found that ∼50% of human tissues present a positive correlation between stemness and proliferation and 20% a negative correlation. Furthermore, all our analyses show negative correlations between stemness and cellular senescence, with significant results in ∼80% of the tissues analyzed. Finally, we also observed a trend that hematopoietic stem cells derived from old patients might have more stemness. In short, we assigned stemness scores to human samples and show evidence of a pan-tissue loss of stemness during human aging, which adds weight to the idea that stem cell deterioration contributes to human ageing.

## Main text

Although the aging process is the leading cause of human mortality and morbidity associated with several diseases, scientists still debate its causes and mechanisms (1, 2). Among the biological pathways associated with aging, we can highlight stem cell exhaustion, which basically describes that during normal aging, the decrease in the number or activity of these cells contributes to physiological dysfunctions in aged tissues (3). This concept is reinforced by the fact that aging is associated with reduced tissue renewal and repair at advanced ages (4). Moreover, longevity manipulations in mice often affect growth and cell division, which has been hypothesized to relate to stem cells (5).

Despite its importance, *in vivo* detection and quantification of stem cells is still challenging, which makes it difficult to study its association with aging, especially in humans (6). In this context, detecting stemness-associated expression signatures is a promising strategy for studying stem cell biology. Stemness is defined as the molecular processes that characterize the fundamental properties of stem cells for the generation of daughter cells and self-renewal, and although it is a concept widely used in oncology, it has been little studied in gerontology. (7, 8, 9).

In this study, we applied a machine learning method to detect stemness signatures from transcriptome data of healthy human tissues. To do this, we first downloaded expression data of 17,382 samples, divided into 30 tissues aged between 20 and 79 years, from GTEx in transcripts per million (TPM) (10). After that, we followed step by step the methodology created by Malta et al. and assigned a stemness score to all GTEx samples (8). Briefly, the stemness score varies from 0 (lowest stemness of the samples) to 1 (highest stemness of the samples), and this methodology went through rigorous validation steps by Malta et al., including tests in several datasets from tumor and non-tumor samples. All the data generated are in supplementary table 1 and include the stemness score and clinical data from GTEx.

First, in Figure 1A, we show the distribution of stemness in the 27 human tissues with at least 50 samples. We notice that the highest stemness scores are in testicles and blood, which are known to have a higher number of stem cells (11, 12). We also see that the highest stemness scores are below 0.7, and this is because values above that are in EBV-transformed lymphocytes samples and not present in the graph.

**Figure 1:**
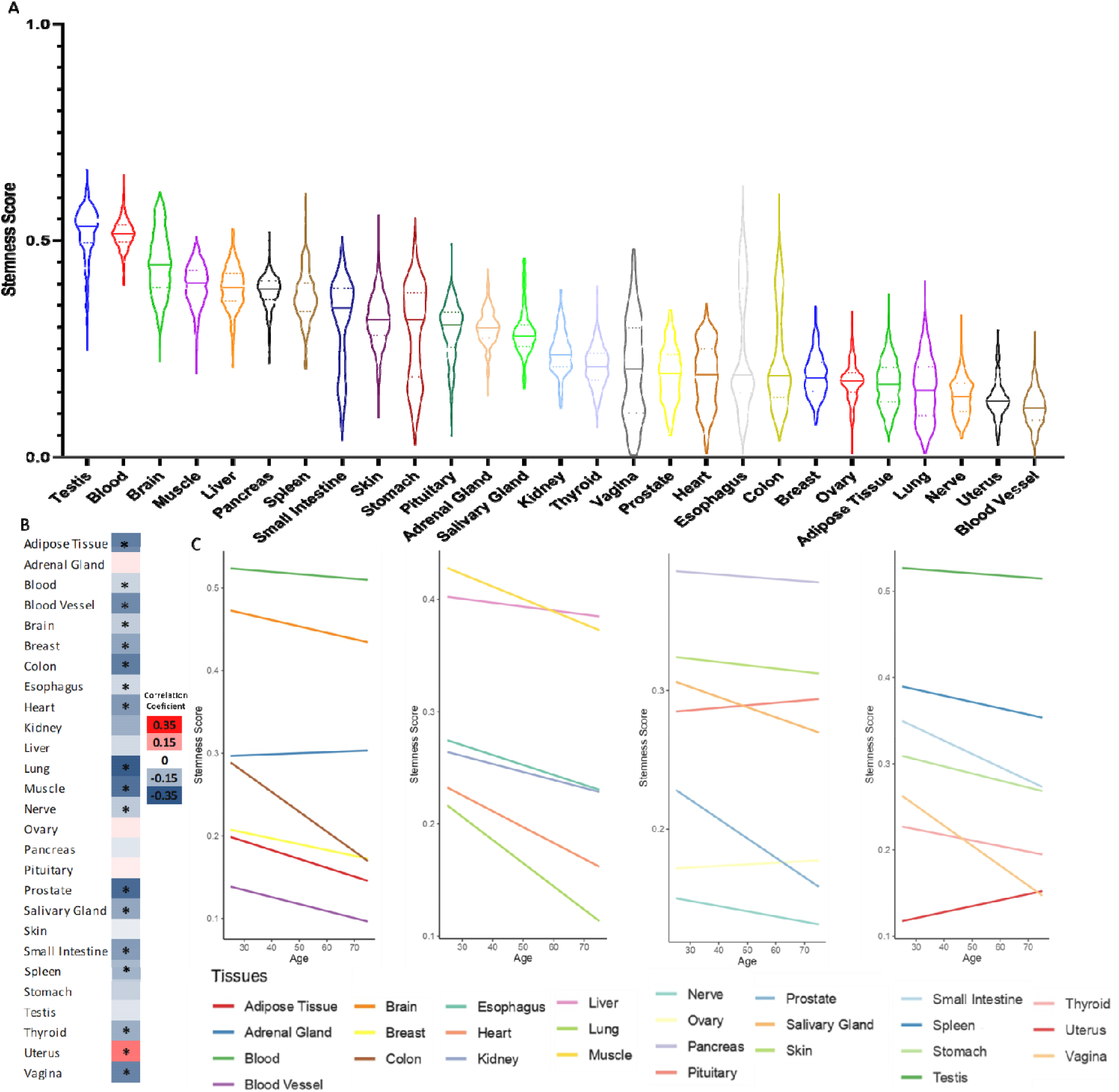
Stemness levels during human aging. **A** Distribution of stemness in human tissues. **B** Heatmap of Pearson’s correlation coefficient between stemness scores and age in human tissues. **C** Linear trend between stemness scores and age in human tissues. *FDR <0.05

Then, we analyzed the relationship between the stemness and aging. In Figure 1B, we have a heatmap of the correlation coefficient (Pearson correlation test followed by Benjamini-Hochberg correction) between stemness score and age, and in Figure 1C, we show the linear trend of the same variables. We observe a pan-tissue loss of stemness in most tissues studied, with the only significant exception being the uterus. Interestingly, our group previously showed that the uterus also tends to behave differently concerning cellular senescence and gene expression patterns during aging (13, 14).

To explore potential mechanisms, we associated stemness with cell proliferation and senescence, two processes associated with aging. We investigated the association between stemness and cell proliferation by downloading the gene expression (in TPM) of the proliferation marker MKI67 and correlated it with the stemness score of the samples. Figure 2A shows the correlation between stemness and proliferation in all GTEx samples (Pearson’s correlation test). Figure 2B shows a heatmap of the correlation coefficient in all tissues from GTEx (Pearson’s correlation test followed by Benjamini-Hochberg correction).

**Figure 2:**
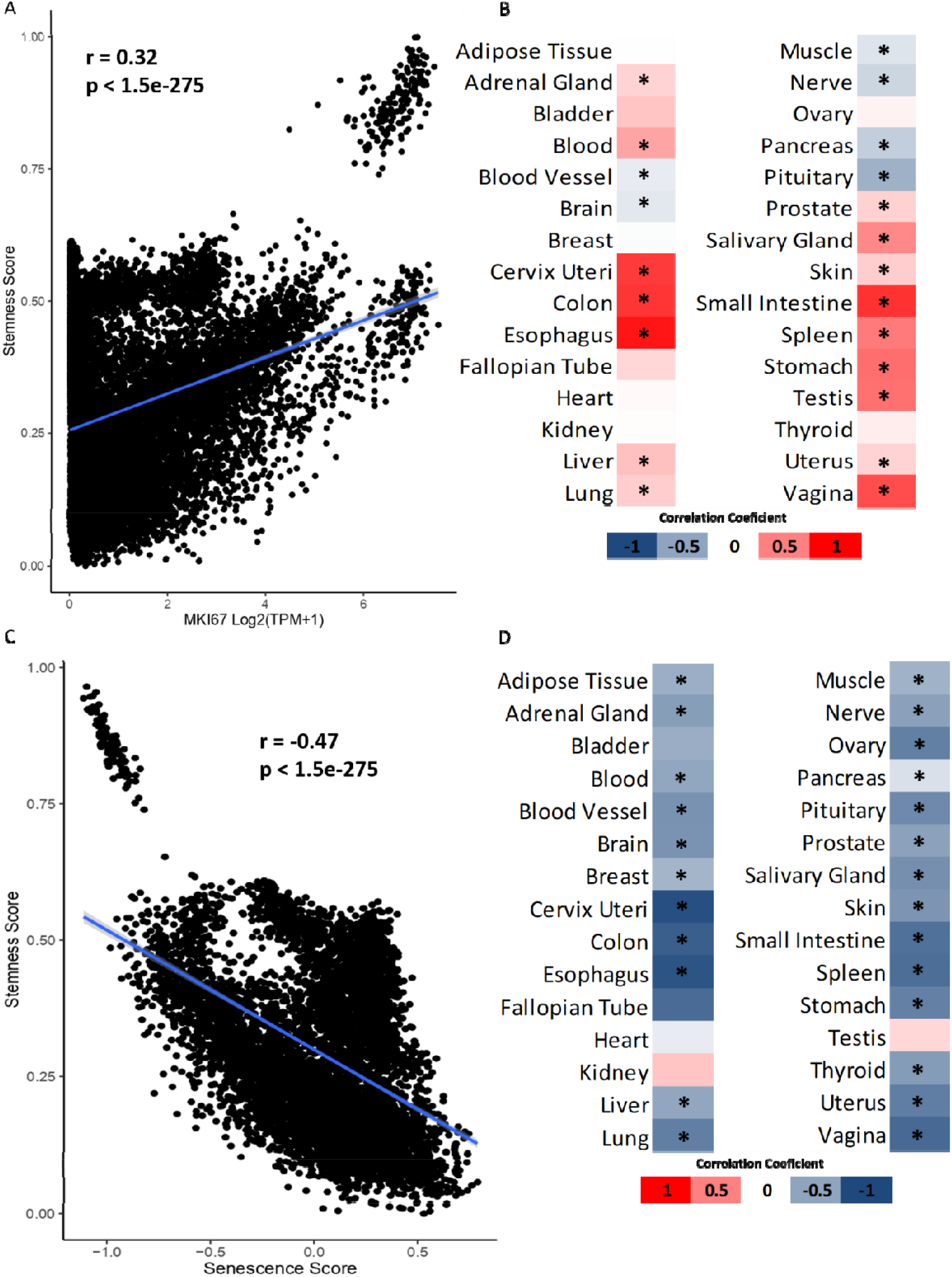
Relationship between stemness, cellular proliferation and senescence. **A** Correlation between stemness score and MKI67 expression. **B** Heatmap of Pearson’s correlation coefficient between stemness scores and MKI67 expression. **C** Correlation between stemness score and senescence scores. **D** Heatmap of Pearson’s correlation coefficient between stemness score and senescence score. *FDR <0.05

We clearly observe a general positive correlation trend between stemness and proliferation, but this correlation is not observed in all tissues as exceptions include blood vessel, brain, muscle, nerve, pancreas, and pituitary results (Figure 2 A-B). This suggests that, although cell proliferation is important for stemness, their relationship is more complex and may not be as straightforward.

Then, we used Wang et al. senescence score data to verify the association between stemness and cellular senescence (15). In total, we have 7123 samples with stemness and senescence values simultaneously, and the correlations were performed as before. We can observe an opposite correlation between stemness and senescence when considering all available GTEx samples (Figure 2C) and when we separate by tissue (Figure 2D) without any significant exception. That is interesting because it indicates that although senescent cells and stem cells are not technically opposite states, they behave in opposite ways *in vivo* at the transcriptomic level.

Finally, since we are working with tissue samples (and therefore analyzing a pool containing somatic and stem cells), we asked ourselves if the stemness of stem cells varies with age. For this, we used data from Adelman et al (16) to compare the stemness levels directly in hematopoietic stem cells (HSC) isolated from humans of two age groups: young (age 18-30) and old (age 65-75).

We observed a quite unexpected result; despite not having a statistically significant difference, we see that both the comparison between the groups (Figure 3A, p=0.07) and the direct correlation (Figure 3B, r=0.42 and p=0.068), have an indicative trend of HSC from older donors to have more stemness. This result is interesting when we consider that literature shows evidence of increasing HSC numbers with age, which may suggest that age-related stem cell problems are functional rather than quantitative (16). Nevertheless, more robust studies must be performed before we can draw more assertive conclusions.

**Figure 3:**
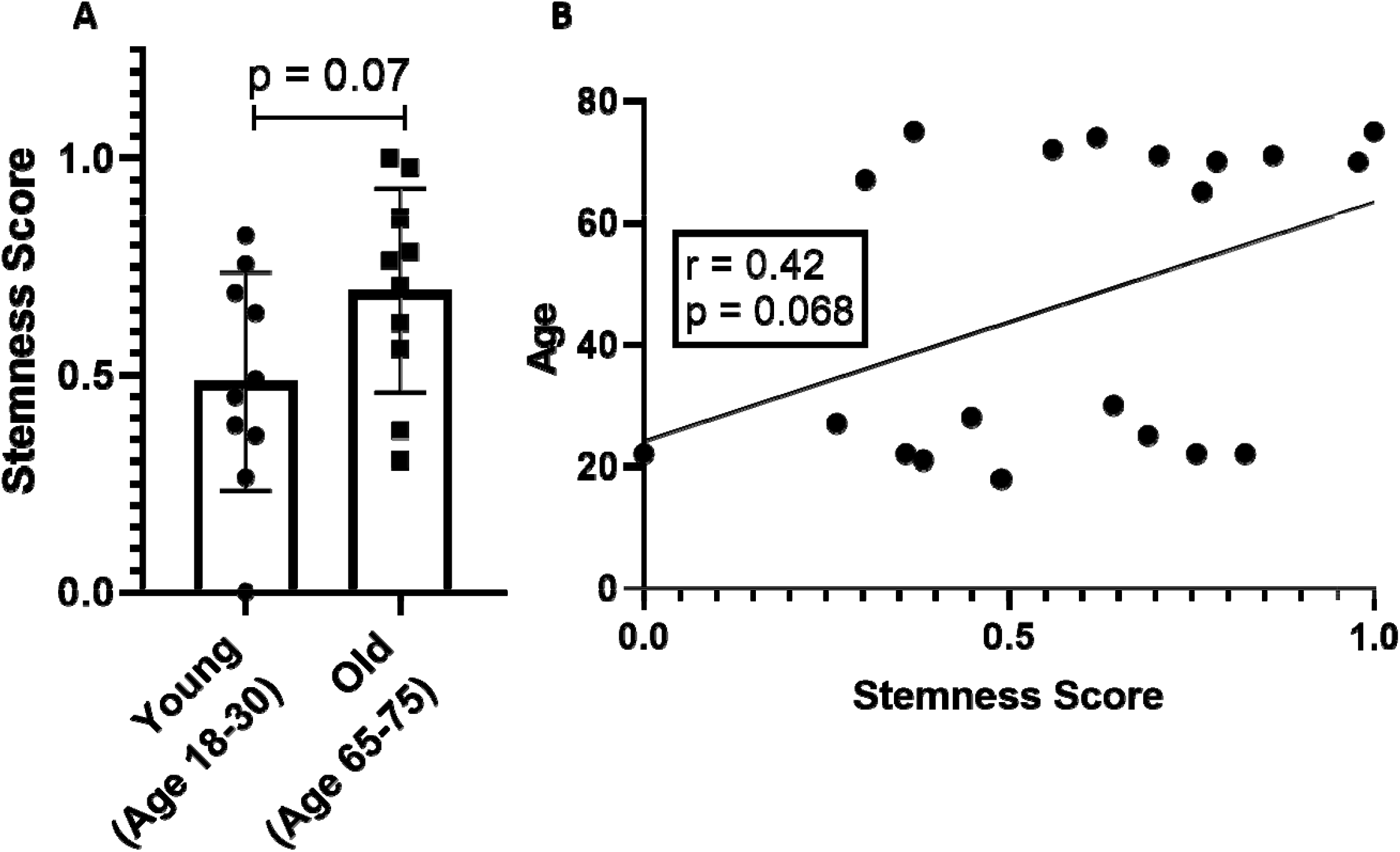
Stemness levels in hematopoietic stem cells. **A** Direct comparison between young and old groups. **B** Correlation between stemness score and age of the HSC donors.

In this report, we studied the relationship between stemness and aging in humans. We found a loss of stemness in most tissues, which corroborates that stem cell exhaustion is important for aging. Although stem cell depletion in aging has already been confirmed in some tissues, such as satellite cells in the muscle (17), and hematopoietic stem cells in blood and bone marrow (18), as far as we know, we were the first to provide evidence of this in a pan-tissue manner.

We also showed the existence of a positive trend between stemness and cell proliferation, but it is not global, with exceptions in 20% of the analyzed tissues. When we consider cellular senescence, we observe that the two phenomena are opposite in all the significant results.

It is important to note that many questions remain open. Our analyses are all correlation tests; therefore, we cannot determine cause and effect. This is important as we cannot assure whether the loss of stemness contributes to aging or is a cause or response of it. Furthermore, we do not know whether the decrease in stemness is a direct reduction of the stem cell pool or refers to intrinsic characteristics of different cells in the tissue. In this sense, our HSC analyses corroborate a functional problem in aged stem cells, at least in blood stem cells. Ultimately, it is crucial to determine what drives these changes and which genes are associated with this process. Here is essential to highlight some papers that suggest that epigenetic modifications regulate stemness and could be a promising area for future studies (19, 20).

In conclusion, we provide the first evidence of a pan-tissue decrease of stemness during human aging, besides the association between stemness and cell proliferation and senescence. This study also assigned a stemness score to more than 17,000 human samples, and these data can be beneficial for the scientific community for further studies.

## Methods

### Transcriptome data from healthy tissues

RNA-Seq-based gene expression data from human tissues were downloaded from the GTEx portal (https://gtexportal.org) in transcripts per million (version 8) (10). The raw RNA-Seq data were previously aligned to the human reference genome GRCH38/hg38 by the GTEx consortium.

We reduced the number of genes from the GTEx data to the same pairs found in the training matrix. The resulting matrix contained 12,471 mRNA expression values and was used to calculate the stemness score. Here it is important to note that GTEx does not provide the actual age of each sample but rather age ranges (20-29, 30-39, 40-49, 50-59, 60-69, and 70-79). We then approximate the age of each sample to 25, 35, 45, 55, 65 and 75 years, respectively, as previously (13).

### Stemness score

To assign the stemness score on the GTEx samples we used a machine learning methodology built by Malta et al. (8).

In brief, the authors built a predictive model using a one-class logistic regression on the pluripotent stem cell samples (ESC and iPSC) from the PCBC dataset (21, 22, 23). The data were mean-centered, and the logistic regression was applied to the stem cell labeled samples to obtain the training signature. We then applied this signature to the GTEx transcriptome data, using Spearman correlations between the model’s weight vector and the sample’s expression profile. As a result, we have a stemness score for all GTEx samples ranging from low (zero) to high (one) stemness. Further details and validation of the methodology can be found in the original study (8).

All this process was done using R and the code available on GitHub associated with the original paper (8). We followed the author’s guidelines step by step.

### Analysis of hematopoietic stem cells

We first downloaded normalized counts of bulk RNA-seq from the work of Adelman et al (16). In this study, the authors isolated human hematopoietic stem cells from two groups with ten samples each: young (age 18-30) and old (age 65-75). We applied the same approach as above to generate stemness scores in these samples. We then compared stemness with age by direct comparison between the two groups (Student’s t-test) and by Pearson correlation. The graphs for this analysis were constructed using GraphPad Prism 8.

### Statistical analysis and graphs

We applied Pearson correlations to all analyses performed in this work using basic R functions. In tissue-specific analyses, the p-value was corrected using Benjamini-Hochberg’s FDR methodology.

Linear trends and the respective graphs were built using the ggplot2 (version 3.3.6) with standards parameters (24). Heatmaps and violin plot were built using Microsoft Excel and GraphPad Prism 8, respectively.

## Acknowledgements

We are grateful to current and past Genomics of Ageing and Rejuvenation Lab members for valuable discussions.

## Notes

**Conflict of interest:** JPM is CSO of YouthBio Therapeutics, an advisor/consultant for the Longevity Vision Fund and NOVOS, and the founder of Magellan Science Ltd, a company providing consulting services in longevity science.

**Funding:** Gabriel Arantes dos Santos is financed by the scholarship “Bolsa de Excelência em Medicina Domingos Nelson Martins” of the Faculty of Medicine of the University of São Paulo (FMUSP). Work in our lab is supported by grants from the Wellcome Trust (208375/Z/17/Z), Longevity Impetus Grants, LongeCity and the Biotechnology and Biological Sciences Research Council (BB/R014949/1 and BB/V010123/1).

### Competing Interest Statement

JPM is CSO of YouthBio Therapeutics, an advisor/consultant for the Longevity Vision Fund and NOVOS, and the founder of Magellan Science Ltd, a company providing consulting services in longevity science.

